# TattleTail: A Pyocin Prediction Tool

**DOI:** 10.64898/2026.03.25.712926

**Authors:** Rayhaan G. Pais, Weilian Chen, Sebastian Leptihn, Xiaoting Hua, Belinda Loh

## Abstract

Tailocins are phage tail-like bacteriocins (PTLBs) thought to be remnants of prophages that have lost the ability to package viral genomes while retaining the ability to kill closely-related bacterial strains, thereby mediating bacterial competition. Tailocins produced by *Pseudomonas aeruginosa* are referred to as pyocins. Apart from their contribution to ecological fitness, they also have the potential to be harnessed as highly-specific antimicrobials to treat antibiotic resistant bacterial infections. Although pyocins lack the genetic components to package viral genomes, pyocin-encoding gene clusters share a high degree of genetic homology to phage tail genes, attributed to their shared ancestry. This poses a significant annotation-based challenge, as current prophage prediction tools, which rely on phage homology for prediction, can misclassify pyocins or tailocins as prophages. Pyocins unknowingly being misannotated as prophages is not only a bioinformatic issue, but can certainly confound experiments examining bacterial competition and prophage induction, if the experimental setup is based on this unintentional misannotation. In this study, we present “TattleTail”, the first version of a bioinformatic tool designed to accurately identify tailocins in genome sequences, with a focus on identifying phage tail-derived pyocin-encoding gene clusters in *P. aeruginosa* in its first iteration. The tool leverages conserved pyocin gene cluster markers and accounts for the absence of canonical phage features, such as capsid, terminase and integrase genes, thereby distinguishing pyocins from intact and cryptic prophages. Validation in *P. aeruginosa* and non-*P. aeruginosa* genomes confirmed the presence of pyocin regions in all *P. aeruginosa* genomes, while none were detected in any non-*P. aeruginosa* genomes. Notably, TattleTail enabled the identification of representative pyocin-encoding gene clusters in clinical *P. aeruginosa* isolates. The identified pyocins in the clinical isolates were induced using mitomycin C, visualized via transmission electron microscopy, processed via tangential flow filtration and demonstrated bactericidal activity, thereby confirming TattleTail predictions. TattleTail aims to complement existing prophage prediction tools during genomic analyses involving phage-derived elements in bacterial genomes, allowing more accurate identification of these elements facilitated by robust discrimination between prophages and tailocins.

## Introduction

Bacteria harbour and employ a variety of defense mechanisms and antagonistic strategies against, but not limited to, closely related bacterial species to compete in densely populated microbial communities. Among these strategies, quiescent bacteriophages, known as prophages, and bacteriocins play pivotal roles in bacterial competition, contributing to the shaping of microbial communities (1,2). Integrated prophages can confer fitness advantages to their bacterial hosts by encoding antibiotic resistance genes, toxins, virulence genes, and by providing superinfection immunity, thereby influencing host metabolism and physiology (3,4). Upon prophage induction, which can be triggered by bacterial stress-associated factors, bacteriophage-mediated lysis against competitor bacterial species can occur, eliminating rival strains (5). In this manner, prophages, in their integrated or induced state, play an important role in shaping microbial communities.

Bacteriocins are antimicrobial peptides, proteins or protein complexes produced by bacteria for a similar competitive reason, i.e., killing or inhibiting competing bacterial strains occupying the same ecological niche (6). Among them, tailocins constitute a class of phage-tail derived bactericidal protein complexes, making them a distinct entity from prophages, but very much evolutionarily related (7,8). Tailocins are thought to be remnants of prophages that have lost the ability to package and transmit viral genomes (7), yet retain the capacity to target and kill closely-related bacterial strains upon induction (9). Consequently, tailocins lack canonical prophage genes, such as capsid proteins, terminases and integrases, but retain several tail fiber proteins and phage-tail-related proteins that mediate killing of competitor strains (8).

Tailocins produced by *Pseudomonas aeruginosa* are specifically referred to as pyocins. Pyocins are broadly classified into three major types: the phage tail-like R- and F-type, and the soluble S-type pyocins (9). R-type and F-type pyocins resemble Myovirus-like and Siphovirus-like phages respectively (10). *P. aeruginosa* strains can harbor a highly conserved pyocin gene cluster that may encode an R-type, an F-type or both R- and F-type pyocins within the same biosynthetic gene cluster (11).

Despite their functional divergence, prophages and pyocins, or more broadly, tailocins, share a substantial amount of genetic and structural similarity due to their common evolutionary origin (12). Although pyocin biosynthetic gene clusters lack canonical phage genes such as capsid, terminase and integrase-related genes, they retain homologues of structural tail proteins, as well as proteins involved in assembly, regulatory factors and lysis (13,8). These shared features strengthen the hypothesis that tailocins are indeed functional prophage remnants. However, these same genomic features also present a challenge for genomic analyses, as they can obscure the accurate distinction between prophages and tailocins in bioinformatic predictions. This ambiguity poses a particular challenge for genome annotation, including prophage prediction and comparative genomics. State-of-the-art prophage prediction tools, such as PHASTEST (14), enable efficient and accurate identification of true and intact prophages, providing information about prophages in a bacterial strain that is crucial for designing experiments related to horizontal gene transfer, prophage induction and bacterial competition studies. PHASTEST is one of the many prophage prediction tools which has revolutionized the field of prophage biology, allowing for easy, web-based annotation of prophages in bacterial genomes (14–17).

However, most prophage prediction tools rely on sequence similarity to known phage proteins (homologs of phage genes) and the presence of “keywords” related to phages, such as “tail” “fiber”, “capsid”, “integrase” etc. Given the substantial overlap in gene content between prophages and tailocins, such approaches could lead to the misannotation of pyocin-gene clusters as prophages. These errors can distort genomic records by inflating estimates of prophage abundance and of HGT rates in bacterial strains, especially in large-scale genomic studies. Misannotation may also confound bacterial competition experiments if a pyocin remains unaccounted for in an experimental setup. Additionally, as another result, the true distribution and diversity of tailocins could be underestimated and underrepresented.

Currently, the identification of pyocin-encoding gene clusters in *P. aeruginosa* and other bacteria relies on manual inspection. This process is a time-consuming and cumbersome process, particularly for poorly annotated or unannotated genomes, making tailocin identification highly impractical in large-scale analyses, such as studies assessing the prevalence and diversity of pyocins across extensive *P. aeruginosa* datasets.

Here, we present “TattleTail”, a bioinformatic tool specifically designed to identify tailocin-encoding biosynthetic gene clusters, with a current focus on pyocins in *P. aeruginosa*. TattleTail was developed to complement existing prophage prediction tools and enable more robust and accurate distinction between prophages and tailocins in bacterial genomes. This tool and its succeeding, improved versions aim to provide a foundation for improved genome annotation, large-scale examination and analysis of tailocins, and the possible discovery of novel tailocin systems, particularly in bacterial strains wherein tailocin biology is understudied.

Owing to their high specificity, pyocins, and more broadly, tailocins, have been proposed as precision antibacterial agents for therapeutic applications (8). Moreover, as protein complexes rather than replicative viruses, such as bacteriophages, they may present fewer regulatory and safety challenges in clinical development. Accordingly, the identification of undiscovered tailocins is important not only for advancing genomic and experimental research, but also for enabling their development as targeted antimicrobial therapies.

## Materials and Methods

### Architecture of TattleTail

In *P. aeruginosa* specifically, the organism of interest for the first version of TattleTail, the presence of a potential pyocin-encoding gene cluster can be identified on the basis of certain evidence markers. i) The presence of the anthranilate synthase gene pair *trpE* and *trpG* flanking a genomic region is indicative of a known pyocin insertion hotspot (18). ii) The absence of the canonical capsid, terminase and integrase genes that are attributed to complete and intact prophages, but not pyocins. iii) The presence of the pyocin-encoding regulatory genes *prtN* (promoter) and *prtR* (repressor). iv) The presence of holin and endolysin genes in the biosynthetic gene cluster. The decision logic of TattleTail was built based on sequential evaluation of these evidence markers. For the most recently updated instructions on how to use the tool, please visit this page (https://github.com/xthua/tattletail).

### Bacterial genomes for validating the functioning of TattleTail

The set of bacterial genomes used to validate the functioning of TattleTail were acquired from the NCBI RefSeq database and Genbank. The set contained 98 *P. aeruginosa* genomes containing pyocin-encoding gene clusters and 19 non-*P. aeruginosa* bacterial genomes (Supplementary Table 1).

### Bacterial strains

The reference strain *P. aeruginosa* ATCC 27853 was acquired from the German Collection of Microorganisms and Cell Cultures GmbH (DSMZ). The three clinical *P. aeruginosa* strains, *P. aeruginosa* 003, *P. aeruginosa* 005 and *P. aeruginosa* 012, as well as the target strain, *P. aeruginosa* 008, for the evaluation of bactericidal activity of pyocins used in this study, are isolates from the University Hospital in Leipzig.

### Whole genome sequencing of clinical isolates

Bacterial genomes of the three clinical *P. aeruginosa* isolates were extracted using the Monarch® Genomic DNA Purification Kit (New England Biolabs) according to the manufacturer’s instructions. Whole genome sequencing was performed via Illumina sequencing. Sequencing files were adapter-trimmed using Trim Galore v0.6.10 program (19). The resulting files were normalized and error-corrected using the BBnorm function of BBMap v38.98 (20). The genomes were assembled using SPAdes v3.15.3 (21). The resulting Genbank file was then uploaded to the Proksee webserver for the creation of a genomic map (22). The genome sequence was then downloaded from the Proksee webserver in its concatenated (single FASTA) format. The whole genome sequence for the reference strain, *P. aeruginosa* ATCC 27853, was already available (RefSeq ID: NZ_CP101912.1). Therefore, genome extraction and whole genome sequencing for this strain was not performed.

### Prediction of prophages in clinical *P. aeruginosa* isolate genomes

The assembled draft genome sequences of the clinical *P. aeruginosa* isolates were uploaded to the PHASTEST Version 3.0 webserver (14) (PHAge Search Tool with Enhanced Sequence Translation) and analyzed using both lite and deep annotation modes. Similarly, the accession number for the whole genome sequence of *P. aeruginosa* ATCC 27853 (NZ_CP101912.1) was uploaded to the PHASTEST webserver and analyzed. Each predicted prophage was then evaluated manually based on the criteria mentioned in the “Architecture of TattleTail” subsection of the Methods in order to confirm whether they could be pyocin-encoding regions instead of true prophages.

### Prediction of pyocin-encoding genes clusters in clinical *P. aeruginosa* isolate genomes via TattleTail

To assess the performance of our pyocin identification tool, TattleTail, we analyzed a reference strain, *P. aeruginosa* ATCC 27853 and three clinical *P. aeruginosa* genomes. Using TattleTail, we processed each genome under the following parameters: “E-value”: 1e-10, “pident”: 35.0, “length”: 50, “bitscore”: 50.0, “window”: 15000, “min_cluster_span”: 13400. The tool successfully identified one candidate pyocin cluster in each isolate, satisfying all four diagnostic criteria below:

1. presence of the flanking gene pair *trpE* and *trpG*.
2. absence of phage-related genes (capsid, terminase, integrase),
3. presence of the regulatory genes *prtN* and *prtR*.
4. presence of the functional lysis genes holin and endolysin.

### Manual determination of pyocin type

The amino acid sequences of the tail fiber proteins of the predicted regions were manually subjected to pairwise alignments (EMBOSS Stretcher) (23) against the tail fiber protein sequences of representative pyocin types: R1 (WHT11591.1) from *P. aeruginosa* S14 (11), R2 (NP_249311.1) from *P. aeruginosa* PAO1 (24), R5 (DBA05840.1) from *P. aeruginosa* isolate 3-5 (10) in order to determine the pyocin type (10). Currently, TattleTail does not determine pyocin type and therefore the pairwise alignment and subsequent pyocin type determination was carried out manually and independent of the tool.

### Induction and tangential flow filtration (TFF) of pyocins identified in clinical *P. aeruginosa* strains

A single colony of each clinical *P. aeruginosa* isolate was grown overnight in 5 mL of Lysogeny Broth (LB) at 37°C, 220 RPM. The resulting overnight cultures were diluted 1:20 (v/v) into 100 mL of LB in 500 mL Erlenmeyer flasks and grown at 37°C, 220 RPM until OD_600_ ≈ 0.60. Pyocin production was induced as previously described ((25) accepted on 03.03.2026 at “Microbiology”). Briefly, mitomycin C was added to a final concentration of 3 μg/mL, followed by incubation at 37°C, 220 RPM for ∼18 h. Cultures were then treated with 10 μL chloroform to lyse residual cells and centrifuged at 8,000 x g for 30 min at 4 °C. Supernatants were subsequently filtered through sterile 0.22 μm filters and stored at 4 °C.

Uninduced controls for each strain were prepared in parallel under identical conditions, except that mitomycin C was omitted. The uninduced cultures were also lysed with 10 μL chloroform prior to centrifugation and filtration.

Small molecule removal and buffer exchange of pyocins from LB to storage buffer (10 mM Tris-HCl, pH 7.0; 10 mM MgSO_4_) were performed by tangential flow filtration (μPulse TFF system) with a 300 kDa MWCO PES filter according to the manufacturers’ instructions. Twenty milliliters of pyocin-containing supernatant were exchanged into 20 mL of storage buffer, and retentates were stored at 4°C. Uninduced control samples were processed identically.

### Transmission electron microscopy

EM sample preparation was performed as previously described ((25) accepted on 03.03.2026 at “Microbiology”). Briefly, pyocin samples were absorbed onto thin carbon films, washed, negatively stained with 2% (w/v) aqueous uranyl acetate solution (pH 5.0) and examined with an EM 910 transmission electron microscope (Carl Zeiss, Oberkochen, Germany) at an acceleration voltage of 80 kV. Size determination of particles was performed using ITEM Software (Olympus Soft Imaging Solutions, Münster).

### Spot test killing assays

A single colony of *P. aeruginosa* 008 was grown in LB at 37°C with shaking at 220 RPM to an OD_600_ of 0.60. Soft-agar overlays were prepared by mixing the culture with 0.7% (w/v) LB soft agar at a 1:10 (v/v) ratio, followed by vortexing and pouring onto LB agar plates. After solidification, 2 μL of serially diluted TFF-processed pyocin preparations (10^0^ to 10^-4^ in storage buffer) were spotted onto the bacterial lawn and allowed to dry. TFF-processed uninduced preparations were diluted and spotted in the same manner. Plates were incubated at 37°C for 16 hours before zones of growth inhibition were recorded. All assays were performed in biological triplicates.

## Results

### Prophage prediction tools may misannotate pyocin gene clusters as intact prophages

We previously reported that the prophage prediction tool PHASTEST misannotated a pyocin biosynthetic gene cluster in a reference *P. aeruginosa* strain as an intact prophage region, despite the absence of capsid and genome-packaging functions ((25) accepted on 03.03.2026 at “Microbiology”). Here, we extend this observation by showing similar misannotations by PHASTEST in three additional *P. aeruginosa* clinical isolates.

Across these genomes, pyocin loci were classified as intact prophages in 3 instances (including the previously reported case), and as a questionable prophage in 1 instance (Supplementary Table 2, detailed output from PHASTEST in supplementary information 1). These results demonstrate that tailocin loci can be conflated with prophages in standard annotation workflows, with potential implications for studies of prophage induction and lysogeny.

This recurring misannotation highlights a limitation of current prophage prediction tools and motivated the development of TattleTail, a dedicated approach for distinguishing pyocin-encoding clusters from prophage regions.

### Development of the TattleTail pyocin-encoding gene cluster prediction pipeline

TattleTail was implemented as a rule-based pipeline that sequentially evaluates conserved genomic markers and the absence of canonical phage functions. The pipeline first screens each genome assembly for regions flanked by the conserved tryptophan biosynthesis genes *trpE* and *trpG*, which define the locus known to harbor pyocin gene clusters.

Within these candidate regions, TattleTail then assesses the presence of phage genes that are required for a functional phage but absent in pyocins such as hallmark genes encoding capsids, terminases and integrases; detection of these genes leads to exclusion of the region, whereas their absence is required for a pyocin call. Candidate regions that pass this step are then evaluated for the SOS-responsive regulatory genes *prtN* and *prtR*, whose co-occurrence is used as an internal consistency check that the cluster is embedded in the canonical pyocin regulatory context. Finally, TattleTail assesses the presence of the lytic module genes holin and endolysin, which are essential for pyocin release. Only regions that satisfy all criteria are classified as pyocin-encoding. This decision framework is summarized in Figure 1. For each genome analysed, TattleTail compiles these decisions into a report that summarizes pyocin candidates at both global and cluster levels. The global summary reports, for each genome, the presence of *trpE*/*trpG* flanks, regulatory genes (*prtN*/*prtR*) and lytic genes (holin/endolysin), and flags capsid-, terminase- or integrase-like genes indicative of prophages rather than a pyocin. Cluster-level outputs detail each candidate pyocin region, including contig identifiers, genomic coordinates of the trimmed “core” cluster, stepwise PASS/FAIL calls for flanks, blockers, regulators and lysis modules, and a final verdict line that labels the region as “tailocin candidate”.

**Figure 1:**
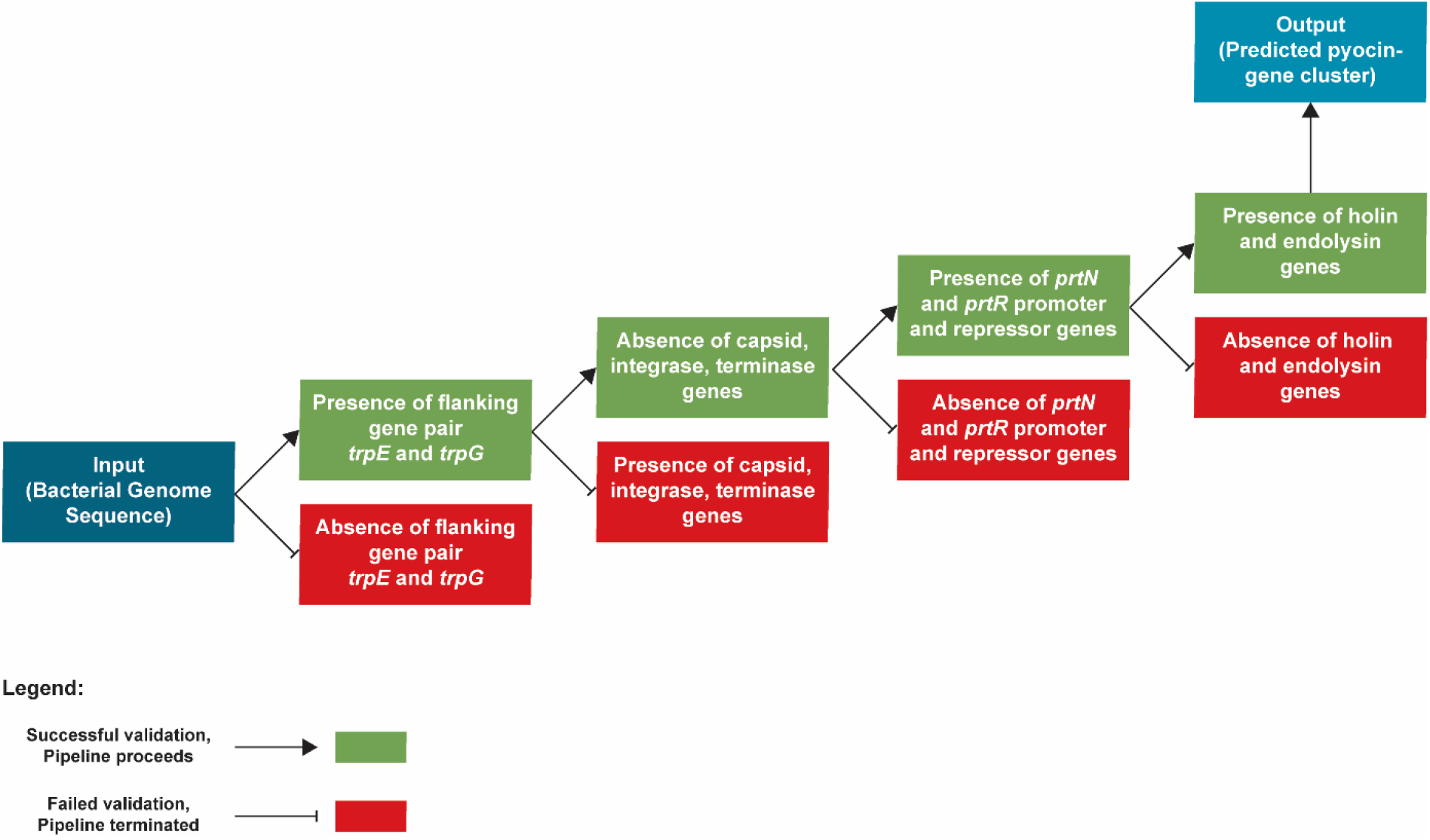
Decision framework of TattleTail Passing branches represent the minimal combination of flanking, regulatory and lytic markers required for a pyocin-encoding gene cluster while explicitly filtering out regions with residual intact phage structural signatures.

In future iterations, we aim to extend the TattleTail GitHub repository with built-in graphical routines that will automatically visualise pyocin loci as fully annotated gene clusters, providing an integrated, interactive view of pyocin predictions across large genome collections.

Collectively, these results confirm that TattleTail reliably identifies pyocin loci in *P. aeruginosa* genomes while avoiding false positives in non-*P. aeruginosa* genomes, addressing the misannotation issues previously observed with prophage prediction tools such as PHASTEST (14), PHAST (26) and ProPhinder (16,27) ((25) accepted on 03.03.2026 at “Microbiology”).

### TattleTail accurately identifies the pyocin-encoding gene clusters in *P. aeruginosa* genomes

To assess the performance of TattleTail, we applied the pipeline to a curated benchmark panel comprising 98 unique *P. aeruginosa* genome assemblies, each containing a pyocin-encoding gene cluster, together with 19 non-*P. aeruginosa* bacterial genomes as negative controls (Supplementary Table 1). Analysis with TattleTail confirmed the presence of pyocin regions in all 98 *P. aeruginosa* genomes, while no pyocin regions were detected in any of the 19 non-*P. aeruginosa* genomes, indicating high sensitivity for pyocin loci in *P. aeruginosa* and high specificity against unrelated bacterial genomes (Supplementary Table 1, detailed run reports in supplementary information 2).

Across the positive genomes, all predicted clusters consistently satisfied all four decision criteria defined in the decision framework. Inspection of TattleTail reports further showed that these predictions were supported by high-confidence homology matches for pyocin-associated genes, including high percentage identities and low e-values for *prtN, prtR*, holin and endolysin across isolates (28,8).

Together, these results demonstrate that the knowledge-driven TattleTail framework reliably identifies pyocin loci in *P. aeruginosa* genomes while avoiding false positives in non-*P. aeruginosa* genomes.

### *In silico* identification of pyocin-encoding gene clusters in clinical *P. aeruginosa* isolates via TattleTail

Having established the performance of TattleTail on the curated benchmark dataset, we next applied the pipeline to a reference *P. aeruginosa* strain (ATCC 27853) and a set of clinical *P. aeruginosa* isolates collected at the Leipzig University Hospital that were not included in the benchmark panel. These clinical isolates comprised strains harboring distinct pyocin types and were therefore used as representative isolates for downstream analysis. Draft genomes were assembled using Proksee and analysed with TattleTail using the same rule set and parameters as used previously.

In all genomes with predicted pyocin loci, TattleTail identified a single candidate region located between the *trpE*–*trpG* genes. These regions satisfied the expected pyocin-defining features including regulatory, structural and accessory components. For representative isolates, the resulting cluster reports contained ordered gene lists spanning the entire pyocin region, encompassing regulatory genes, tail-structure components and accessory genes with their annotated products and genomic coordinates, consistent with the genetic organisation described for R- and F-type pyocins in clinical *P. aeruginosa* collections.

In contrast, PHASTEST identified multiple putative prophage regions in the same genomes, with predicted completeness ranging from ‘questionable’ to ‘intact’, and consistently classified the TattleTail predicted pyocin loci as intact or questionable prophages (Table 1 and Supplementary Table 2).

**Table 1:** TattleTail prediction of pyocin-encoding gene clusters in *P. aeruginosa* genomes.

| Bacterial Strain | Number of Candidate Regions | Region Length | Region Position |
| --- | --- | --- | --- |
| *P. aeruginosa* ATCC 27853 | 1 | 17.13 Kb | 679,269 - 696,399 |
| *P. aeruginosa* 003 | 1 | 17.00 Kb | 6,283,495 - 6,300,498 |
| *P. aeruginosa* 005 | 1 | 17.53 Kb | 3,828,968 - 3,846,494 |
| *P. aeruginosa* 012 | 1 | 31.02 Kb | 403,320 - 434,337 |

These observations highlight the difficulty of distinguishing tailocin loci from prophages using general-purpose annotation pipelines. In contrast, TattleTail consistently identified these loci as pyocin-encoding gene clusters without introducing additional prophage-like predictions.

### Induction, morphology validation and preliminary activity of pyocins identified in clinical *P. aeruginosa* strains

To validate that the pyocin-encoding loci identified by TattleTail in the Leipzig clinical isolates encode functional tailocins, pyocin production was induced with mitomycin C and analysed for particle morphology and preliminary bactericidal activity (Figure 2A, 2B).

**Figure 2:**
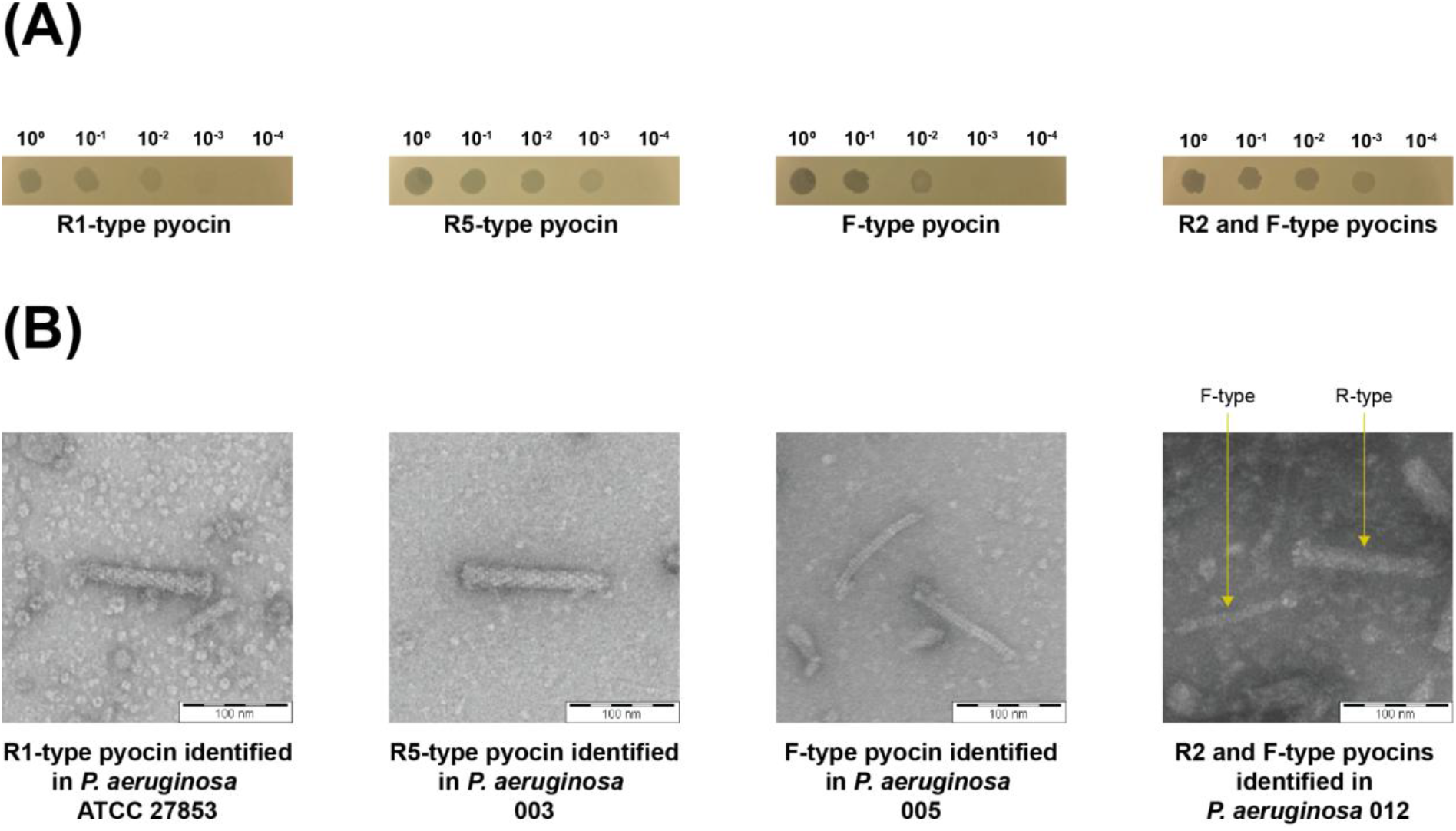
Pyocins identified in clinical *P. aeruginosa* isolates using TattleTail. **(A)** Serial dilution spot test assays of representative pyocin(s) against the target strain *P*. aeruginosa 008. **(B)** Transmission electron micrographs showing the morphologies of each identified and induced pyocin.

Transmission electron microscopy of cell-free supernatants revealed structures characteristic of phage tail-like bacteriocins, including contractile tail particles consistent with R-type pyocins and non-contractile tail particles consistent with F-type pyocins (Figure 2B). No phage heads or DNA-containing capsids were observed, consistent with the absence of capsid and terminase genes in the corresponding genomic loci and supporting the absence of intact prophages.

Activity assays using spot tests on susceptible *P. aeruginosa* indicator strains showed clear zones of growth inhibition for supernatants from induced cultures (Figure 2A), whereas non-induced controls showed significantly less killing (Supplementary Figure 1), suggesting low-level, mitomycin c-independent basal induction.

Additional validation experiments, such as Proteinase K treatment, assessment of self-susceptibility, and evaluation of lack of amplification potential, remain to be performed to further confirm that the observed bactericidal activity is attributable solely to pyocins.

Together, these results demonstrate that the pyocin loci predicted by TattleTail correspond to expressed, bactericidal tailocins in clinical *P. aeruginosa* isolates, providing experimental support for the accuracy of the *in silico* predictions and the validity of the tool.

## Discussion

A wide range of specialised tools are available for the systematic identification of non-essential, accessory elements in bacterial genomes, including antibiotic resistance genes (29,30), integrons (29), transposable elements and insertion sequences (31), pdif-flanked resistance modules (32), group II introns (33) and prophages (15). However, despite the increasing recognition of tailocins as important mediators of bacterial competition, no dedicated genome mining tool has been developed for their identification. TattleTail addresses this gap by providing a targeted bioinformatic approach for the identification of pyocin-encoding gene clusters. In its current implementation, the tool integrates the detection of conserved phage-tail-associated genes, with the exclusion of bona fide whole phage genes, such as capsid genes. This design enables discrimination between pyocin loci and intact prophages, addressing a key limitation in the annotation of phage-derived elements in bacterial genomes.

This capacity is especially advantageous in large-scale genomic studies, where manual inspection and annotation of genomes to uncover pyocin-encoding gene clusters is impractical. For example, systematic re-analysis of genomes in the *P. aeruginosa* database using TattleTail may help clarify whether pyocin-encoding regions are underreported and prophage-encoding regions are correspondingly overestimated. For genomic analyses, when utilized in parallel with tools specialized for accurate prophage prediction, such as PHASTEST, the overall accuracy of a certain genomic analysis will improve in accuracy. Therefore, importantly, TattleTail is not intended to replace existing prophage prediction tools, but rather aims to complement them, enabling more accurate and comprehensive annotation when used in combination.

Accurate genome annotation is critical for experimental design, particularly in studies of bacterial competition, horizontal gene transfer and prophage induction. If such experiments are designed based on misannotated prophages, the results could be misrepresented and confounded, making interpretation difficult. In such contexts, the combined use of TattleTail with established tools for other mobile genetic elements can improve experimental design, and downstream data interpretation.

TattleTail is designed to be scalable and accessible, requiring only genome assemblies as an input. Its structured output facilitates straightforward interpretation, and batch processing makes it practical for analyzing many genomes at once.

When tested on the training set, TattleTail accurately identified pyocin-encoding gene clusters in all 98 *P. aeruginosa genomes*, while maintaining high specificity against non-*P. aeruginosa* genomes, demonstrating its reliability. Application to clinical *P. aeruginosa* isolates further confirmed that the tool robustly detects pyocin clusters in real-world datasets., making it a useful complement to existing prophage prediction tools.

Several avenues exist for further development. Future versions of TattleTail (TattleTail 2.0) will include more in-depth identification and annotation capabilities, including the annotation of all pyocin-associated genes in an identified cluster, classification into R- and F-type systems, and subtype-level resolution (e.g. R1, R2, R5, F8, etc.). Implementation of TattleTail as a web server-based platform is also planned to improve accessibility and usability.

The long-term goal of TattleTail is to expand its identification capabilities beyond pyocins and identify tailocins across diverse bacterial taxa, especially in systems where tailocin biology remains understudied. Such efforts may facilitate the discovery of previously uncharacterized bacterial defense mechanisms, thereby paving the way for novel research in bacterial competition, ecological niche-based studies and support the development of new antibacterial agents. Given their specificity and non-replicative nature, tailocins represent promising candidates for targeted therapeutic applications, particularly in the context of antibiotic resistant infections.

TattleTail is freely provided as a resource for non-commercial use (https://github.com/xthua/tattletail). Its application to *P. aeruginosa* genomes from publicly available databases, and all feedback received will support continued refinement and expansion of its capabilities.

## Authors’ Contributions

RGP, BL and SL conceived the study and designed the decision logic of the TattleTail prediction pipeline. WC and XH implemented the TattleTail code and performed the bioinformatic analyses. RGP carried out the experimental work on clinical isolates. RGP, BL and SL interpreted the results, designed the figures and wrote the manuscript. RGP, BL, SL, WC and XH edited and revised the manuscript critically and all authors approved the final version.

## Acknowledgements

We thank Dr. Mathias Müsken for carrying out transmission electron microscopy. We also thank Vyshnavi Ramireddy for processing (induction, TFF) pyocin samples.

## Data and Code Availability

The software source code, example datasets and documentation are available at (https://github.com/xthua/tattletail) and will be archived at the time of publication under a permanent DOI. The custom BLAST protein database we used in this study can be found in Supplementary information 3. Genome assemblies analysed for the benchmark test in this study were obtained from NCBI RefSeq and GenBank (accession numbers listed in Supplementary Table 1), and genome sequences of clinical *P. aeruginosa* genomes will be deposited in a publicly available genome database. The submission of the genome sequences of the clinical *P. aeruginosa* strains are being processed and are currently not publicly accessible; thus, the authors will provide them upon request.

## Conflict of Interest

Weilian Chen is employed by Hangzhou Digital-Micro Biotech Co., Ltd., Hangzhou, China. The company had no influence on the study design, data analysis, or decision to publish this manuscript.

## Notes

https://github.com/xthua/tattletail

