## Supplementary Table 1 for "TattleTail: A Pyocin Prediction Tool"

**Supplementary Table 1:** List of *Pseudomonas aeruginosa* and non-*Pseudomonas aeruginosa* genomes acquired from the RefSeq and Genbank databases for the validation of the functionality of TattleTail. Region Length and Region Position were determined using TattleTail.

**98 *P. aeruginosa* genomes with pyocin-encoding gene clusters**

| # | Genbank/Refseq | Assembly | Region Length | Region Position |
|---|---|---|---|---|
| 1 | NC_002516.2 | GCF_000006765.1 | 31.02 Kb | 672,459 - 703,476 |
| 2 | CP000438.1 | GCF_000014625.1 | 29.18 Kb | 685,281 - 714,460 |
| 3 | FM209186.1 | GCF_000026645.1 | 17.13 Kb | 663,604 - 680,732 |
| 4 | CP002496.1 | GCF_000226155.1 | 17.14 Kb | 663,464 - 680,599 |
| 5 | AP012280.1 | GCF_000284555.1 | 17.00 Kb | 5,938,573 - 5,955,576 |
| 6 | DF126593.1 | GCF_000291745.1 | 17.11 Kb | 4,622,486 - 4,639,594 |
| 7 | CP006705.1 | GCF_000468555.2 | 31.02 Kb | 672,336 - 703,353 |
| 8 | CP006728.1 | GCF_000468935.2 | 30.97 Kb | 672,436 - 703,406 |
| 9 | CP006831.1 | GCF_000484495.2 | 31.02 Kb | 672,459 - 703,476 |
| 10 | CP006832.1 | GCF_000484545.2 | 31.02 Kb | 672,459 - 703,476 |
| 11 | CP007224.1 | GCF_000626655.2 | 17.22 Kb | 664,485 - 681,700 |
| 12 | KN050641.1 | GCF_000743405.1 | 17.13 Kb | 4,268,061 - 4,285,191 |
| 13 | CP021380.2 | GCF_000763245.3 | 31.02 Kb | 677,714 - 708,731 |
| 14 | AP014646.1 | GCF_000829255.1 | 17.00 Kb | 667,450 - 684,453 |
| 15 | AP014622.1 | GCF_000829275.1 | 17.00 Kb | 667,447 - 684,450 |
| 16 | CVVB01000099.1 | GCF_001180485.1 | 17.00 Kb | 68,961 - 85,964 |
| 17 | CVXP01000055.1 | GCF_001180545.1 | 17.00 Kb | 68,974 - 85,977 |
| 18 | CVVW01000239.1 | GCF_001180705.1 | 17.00 Kb | 106,158 - 123,161 |
| 19 | CVVY01000482.1 | GCF_001180765.1 | 17.00 Kb | 7,202 - 24,205 |
| 20 | CVVU01000111.1 | GCF_001180785.1 | 17.00 Kb | 18,151 - 35,154 |
| 21 | AP014651.1 | GCF_001547955.1 | 17.00 Kb | 753,655 - 770,659 |
| 22 | AP014839.2 | GCF_00154813 | 18.24 Kb | 702,042 - 720,278 |

| | | | | |
|---|---|---|---|---|
| | | 5.1 | | |
| 23 | LODH01000055.1 | GCF_00156086 5.1 | 17.14 Kb | 18,206 - 35,341 |
| 24 | LOHI01000016.1 | GCF_00160158 5.1 | 28.91 Kb | 18,108 - 47,020 |
| 25 | LOHJ01000009.1 | GCF_00160166 5.1 | 29.26 Kb | 378,943 - 408,202 |
| 26 | LOHH01000012.1 | GCF_00160174 5.1 | 17.14 Kb | 18,193 - 35,327 |
| 27 | CP014948.1 | GCF_00160604 5.1 | 29.43 Kb | 663,105 - 692,536 |
| 28 | CP015001.1 | GCF_00179283 5.1 | 31.02 Kb | 677,765 - 708,782 |
| 29 | CP015002.1 | GCF_00179285 5.1 | 31.02 Kb | 708,484 - 739,501 |
| 30 | CP015003.1 | GCF_00179287 5.1 | 31.02 Kb | 677,758 - 708,775 |
| 31 | LVWC01000001.1 | GCF_00180273 5.1 | 31.02 Kb | 4,101,523 - 4,132,540 |
| 32 | LVXB01000001.1 | GCF_00180650 5.1 | 31.02 Kb | 677,817 - 708,834 |
| 33 | CP014999 | GCF_00187026 5.1 | 31.02 Kb | 708,490 - 739,507 |
| 34 | CP013113.1 | GCF_00187952 5.1 | 17.13 Kb | 695,743 - 712,872 |
| 35 | CP013477.1 | GCF_00190019 5.1 | 17.53 Kb | 681,690 - 699,216 |
| 36 | CP013478.1 | GCF_00190022 5.1 | 17.53 Kb | 681,690 - 699,216 |
| 37 | CP013479.1 | GCF_00190026 5.1 | 17.53 Kb | 681,690 - 699,216 |
| 38 | CP020603.1 | GCF_00208575 5.1 | 17.00 Kb | 667,406 - 684,409 |
| 39 | PVXJ01000005.1 | GCF_00303239 5.1 | 17.13 Kb | 50,307 - 67,435 |
| 40 | CP029605.1 | GCF_00343323 5.1 | 17.00 Kb | 707,622 - 724,625 |
| 41 | CP012901.1 | GCF_00357150 5.1 | 17.14 Kb | 6,207,033 - 6,224,167 |
| 42 | CP034429.1 | GCF_00395782 5.1 | 31.02 Kb | 671,976 - 702,993 |
| 43 | CP034430.1 | GCF_00400049 5.1 | 17.00 Kb | 815,560 - 832,564 |
| 44 | CP037925.1 | GCF_00435512 5.1 | 17.13 Kb | 664,894 - 682,026 |
| 45 | CP037926.1 | GCF_00435514 5.1 | 17.13 Kb | 664,827 - 681,959 |
| 46 | CP041354.1 | GCF_00697178 5.1 | 17.13 Kb | 653,236 - 670,363 |
| 47 | CP043483.1 | GCF_00833098 5.1 | 17.00 Kb | 710,592 - 727,595 |
| 48 | CP043549.1 | GCF_00837008 5.1 | 17.00 Kb | 5,614,848 - 5,631,851 |

| | | | | |
|---|---|---|---|---|
| 49 | CP045739.1 | GCF_00966231 5.1 | 17.00 Kb | 759,584 - 776,587 |
| 50 | CP046069.1 | GCF_00967688 5.1 | 18.89 Kb | 674,281 - 693,168 |
| 51 | CP049161.1 | GCF_01104537 5.1 | 17.66 Kb | 669,102 - 686,761 |
| 52 | CP051766.1 | GCF_01311491 5.1 | 17.00 Kb | 6,139,478 - 6,156,482 |
| 53 | CP051768.1 | GCF_01311493 5.1 | 17.00 Kb | 6,093,905 - 6,110,909 |
| 54 | CP051770.1 | GCF_01311495 5.1 | 17.00 Kb | 6,130,550 - 6,147,554 |
| 55 | AP024513.1 | GCF_01840936 5.1 | 29.43 Kb | 698,481 - 727,911 |
| 56 | BSAL01000015.1 | GCF_02601265 5.1 | 29.16 Kb | 18,374 - 47,530 |
| 57 | BSAN01000013.1 | GCF_02601269 5.1 | 29.16 Kb | 110,627 - 139,782 |
| 58 | BSAP01000005.1 | GCF_02601273 5.1 | 29.16 Kb | 238,074 - 267,229 |
| 59 | BSAQ01000003.1 | GCF_02601275 5.1 | 31.02 Kb | 198,596 - 229,613 |
| 60 | BSAR01000003.1 | GCF_02601277 5.1 | 31.02 Kb | 244,746 - 275,763 |
| 61 | BSAT01000022.1 | GCF_02601281 5.1 | 17.00 Kb | 64,525 - 81,529 |
| 62 | BSAU01000004.1 | GCF_02601283 5.1 | 31.02 Kb | 250,828 - 281,845 |
| 63 | BSAW01000005.1 | GCF_02601287 5.1 | 17.00 Kb | 18,510 - 35,513 |
| 64 | BSAX01000005.1 | GCF_02601289 5.1 | 29.26 Kb | 18,403 - 47,662 |
| 65 | BSAZ01000031.1 | GCF_02601293 5.1 | 17.00 Kb | 43,560 - 60,564 |
| 66 | BSBB01000004.1 | GCF_02601297 5.1 | 31.02 Kb | 18,316 - 49,333 |
| 67 | BSBC01000003.1 | GCF_02601299 5.1 | 29.26 Kb | 18,403 - 47,662 |
| 68 | BSBD01000032.1 | GCF_02601301 5.1 | 16.99 Kb | 42,852 - 59,845 |
| 69 | CP123953.1 | GCF_02991682 5.1 | 29.13 Kb | 697,412 - 726,541 |
| 70 | CP136598.1 | GCF_03284738 5.1 | 17.00 Kb | 741,442 - 758,445 |
| 71 | BAABSN010000004 .1 | GCF_03954471 5.1 | 17.00 Kb | 18,377 - 35,376 |
| 72 | CP173019.1 | GCF_04475898 5.1 | 31.02 Kb | 1,675,859 - 1,706,876 |
| 73 | CP173020.1 | GCF_04475899 5.1 | 31.02 Kb | 4,180,820 - 4,211,839 |
| 74 | CP173021.1 | GCF_04475900 5.1 | 17.00 Kb | 670,460 - 687,462 |

| | | | | |
|---|---|---|---|---|
| 75 | CP173022.1 | GCF_04475901 5.1 | 17.00 Kb | 670,460 - 687,462 |
| 76 | CP173023.1 | GCF_04475902 5.1 | 17.00 Kb | 670,435 - 687,437 |
| 77 | CP069363.1 | GCF_04515981 5.1 | 17.43 Kb | 674,197 - 691,621 |
| 78 | LN870292.1 | GCF_90006902 5.1 | 17.53 Kb | 682,172 - 699,698 |
| 79 | LN871187.1 | GCF_90007037 5.1 | 31.02 Kb | 672,464 - 703,481 |
| 80 | LT608330.1 | GCF_90009580 5.1 | 29.18 Kb | 685,282 - 714,461 |
| 81 | LT673656.1 | GCF_90014928 5.1 | 17.49 Kb | 662,390 - 679,877 |
| 82 | LT969520.1 | GCF_90024335 5.1 | 17.00 Kb | 724,783 - 741,786 |
| 83 | UWWR01000001.1 | GCF_90060791 5.1 | 18.89 Kb | 989,547 - 1,008,432 |
| 84 | UWXD01000001.1 | GCF_90060803 5.1 | 17.13 Kb | 636,493 - 653,622 |
| 85 | UWXL01000001.1 | GCF_90060812 5.1 | 18.89 Kb | 2,778,154 - 2,797,044 |
| 86 | LR130527.1 | GCF_90061825 5.1 | 17.14 Kb | 3,787,906 - 3,805,041 |
| 87 | LR130534.1 | GCF_90061828 5.1 | 17.14 Kb | 742,835 - 759,970 |
| 88 | LR130533.1 | GCF_90061830 5.1 | 17.14 Kb | 4,061,846 - 4,078,981 |
| 89 | LR130536.1 | GCF_90061831 5.1 | 17.14 Kb | 673,408 - 690,543 |
| 90 | LR130535.1 | GCF_90061832 5.1 | 17.14 Kb | 673,424 - 690,559 |
| 91 | LR130537.1 | GCF_90061833 5.1 | 17.14 Kb | 673,395 - 690,530 |
| 92 | OX638701.1 | GCF_95169136 5.1 | 29.29 Kb | 1,944,236 - 1,973,520 |
| 93 | CATOQB01000000 3.1 | GCF_95180237 5.1 | 17.00 Kb | 18,091 - 35,093 |
| 94 | OX638610.1 | GCF_95180237 5.2 | 17.00 Kb | 18,091 - 35,093 |
| 95 | CATORO01000000 1.1 | GCF_95180527 5.1 | 17.00 Kb | 674,279 - 691,281 |
| 96 | OX638564.1 | GCF_95180527 5.2 | 17.00 Kb | 674,279 - 691,281 |
| 97 | CATODC01000000 1.1 | GCA_95169136 5.1 | 29.29 Kb | 1,944,236 - 1,973,520 |
| 98 | LR700248.1 | LR700248.1 | 18.89 Kb | 4,594,995 - 4,613,882 |

**19 non-*P. aeruginosa* genomes**

| # | Genbank/Refseq | Assembly | Region Length | Region Position |
|---|---|---|---|---|
| 1 | CM000725.1 | GCA_000161095.1 | - | - |
| 2 | CM000758.1 | GCF_000161715.1 | - | - |
| 3 | AM920437.1 | GCA_000226395.1 | - | - |
| 4 | JACVVK020000003.1 | GCA_018292915.2 | - | - |
| 5 | JAVFKD010000014.1 | GCA_036429605.1 | - | - |
| 6 | JAQQWG010000004.1 | GCA_038362765.1 | - | - |
| 7 | BX950851.1 | GCF_000011605.1 | - | - |
| 8 | CM000748.1 | GCF_000161515.1 | - | - |
| 9 | CM000752.1 | GCF_000161595.1 | - | - |
| 10 | CM000753.1 | GCF_000161615.1 | - | - |
| 11 | CM000756.1 | GCF_000161675.1 | - | - |
| 12 | FO834906.1 | GCF_000968155.1 | - | - |
| 13 | CP015281.1 | GCF_001620335.1 | - | - |
| 14 | AP019314.1 | GCF_003945305.1 | - | - |
| 15 | CP098234.1 | GCF_029224125.1 | - | - |
| 16 | AP035801.1 | GCF_042847795.1 | - | - |
| 17 | FBYD01000004.1 | GCF_900041715.1 | - | - |
| 18 | LT599825.1 | GCF_900093485.1 | - | - |
| 19 | LR586051.1 | GCA_901538445.1 | - | - |
