## Supplementary Table 2 for "TattleTail: A Pyocin Prediction Tool"

**Supplementary Table 2:** PHASTEST prediction of prophage regions in the genome of the reference strain *P. aeruginosa* ATCC 27853 and clinical *P. aeruginosa* isolate draft genomes.

| Bacterial Strain | Region | Region Length | Completeness | Score | Region position |
|---|---|---|---|---|---|
| *P. aeruginosa* ATCC 27853 | 1* | 18.4 Kb | Intact | 130 | 679,586 - 698,056 |
| | 2 | 40.4 Kb | Intact | 114 | 796,824 - 837,266 |
| | 3 | 42.9 Kb | Intact | 110 | 1,337,187 - 1,380,105 |
| | 4 | 55.2 Kb | Questionable | 84 | 2,508,737 - 2,564,035 |
| | 5 | 49.9 Kb | Intact | 150 | 5,102,870 - 5,152,806 |
| | 6 | 15.1 Kb | Questionable | 85 | 5,787,383 - 5,802,527 |
| *P. aeruginosa* 003 | 1 | 57.1 Kb | Intact | 150 | 3,283,954 - 3,341,094 |
| | 2 | 38.3 Kb | Intact | 150 | 6,203,998 - 6,242,311 |
| | 3* | 18.4 Kb | Intact | 140 | 6,283,681 - 6,302,155 |
| *P. aeruginosa* 005 | 1 | 21.4 Kb | Questionable | 90 | 137,662 - 159120 |
| | 2* | 18.9 Kb | Questionable | 80 | 3,827,311 - 3,846,308 |
| *P. aeruginosa* 012 | 1* | 35 Kb | Intact | 150 | 401,663 - 436,707 |
| | 2 | 45.1 Kb | Intact | 110 | 2,442,251 - 2,483,375 |

* Manual inspection during this study revealed that these regions fulfilled conserved criteria of pyocin-encoding gene clusters in *P. aeruginosa* and not that of prophage-encoding regions.
