## Supplementary figures and images for "TattleTail: A Pyocin Prediction Tool"

### Supplementary figure 1

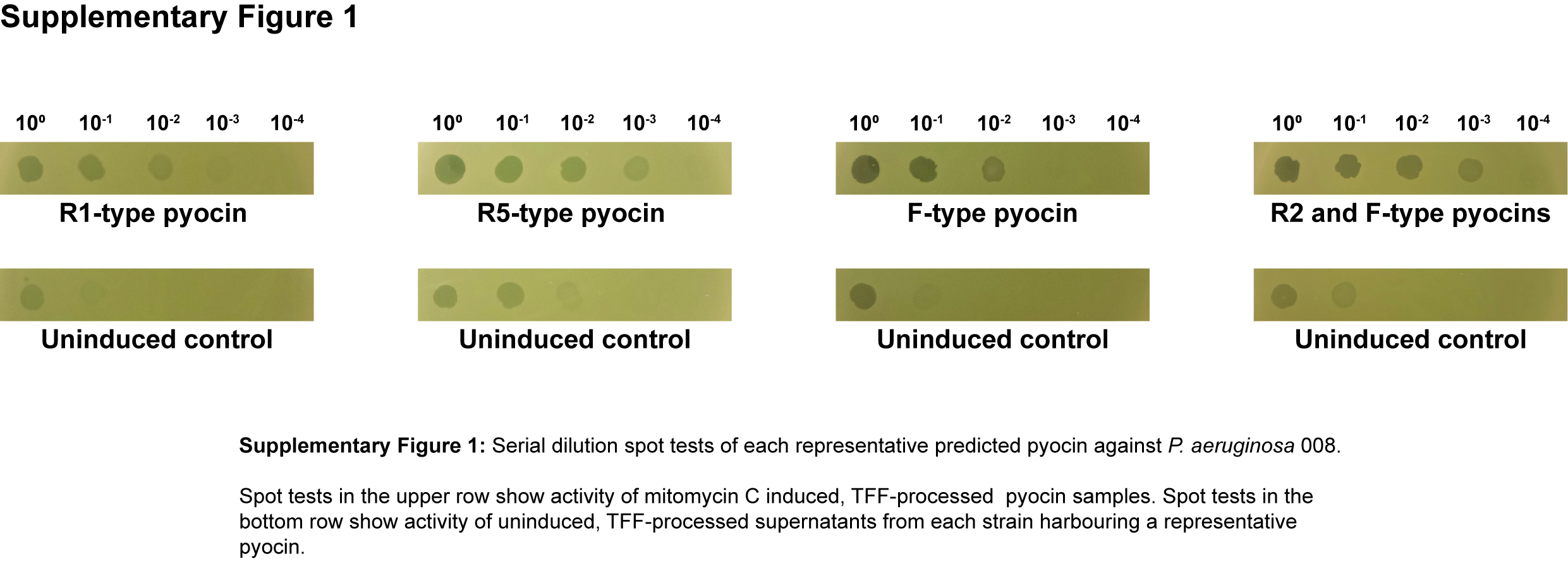
